# HIF isoforms contribute distinctly during human NK cell activation

**DOI:** 10.1101/2025.06.27.661904

**Authors:** Jorrit De Waele, Tias Verhezen, Ho Wa Lau, Jöran Melis, Louize Brants, Jonas Van Audenaerde, Julie de Beukelaar, Charlotte Gevaerts, Filip Lardon, An Wouters, Evelien Smits

## Abstract

Hypoxia-inducible factors (HIFs) are critical transcriptional regulators that enable cellular adaptation to low oxygen environments. Natural killer (NK) cells are key effectors of innate immunity which frequently operate in hypoxic tissues during viral infection and tumor surveillance. In addition, HIF also play a role in NK cells beyond hypoxic adaptation. However, the distinct and overlapping roles of HIF-1α and HIF-2α in NK cell biology remain incompletely understood. In this study, we investigated the contribution of HIF-1α and HIF-2α to NK cell effector functioning in normoxia using pharmacological inhibition. When HIF-1α was inhibited, we observed a pronounced decline in cytotoxicity as well as IFNγ and TNFα production, but not granzyme B, accompanied by elevated mitochondrial reactive oxygen species. In contrast, HIF-2α did not alter these functions, nor did pan-HIF stabilization. These findings indicate a differential role of HIF-1α and HIF-2α in shaping NK cell responses in normoxia, at least during the acute activation phase as investigated here. Still, more mechanistic and contextual insights into the role of HIF isoforms in NK cells is warranted for future therapeutic applications.

## Introduction

Natural killer (NK) cells are cytotoxic lymphocytes of the innate immune system, essential for the early detection and elimination of virally infected and transformed cells. Unlike adaptive lymphocytes, NK cells do not require prior antigen sensitization or clonal expansion to mediate their effector functions^1^. Their cytotoxic activity is tightly regulated through a balance of activating and inhibitory receptors expressed on the cell surface^2^. Upon engagement with target cells, NK cells release cytolytic granules containing perforin and granzymes and secrete immunoregulatory cytokines such as interferon-gamma (IFNγ) and tumor necrosis factor-alpha (TNFα)^3^. These cytokines modulate both innate and adaptive immune responses and play key roles in antiviral and antitumor immunity^4^.

NK cells operate in diverse physiological and pathological environments, including solid tumors, where the tumor microenvironment (TME) imposes multiple layers of immunosuppressive stress^5^. One of the hallmark features of the TME is hypoxia, a condition of reduced oxygen availability caused by the imbalance between oxygen supply and consumption in rapidly proliferating tumors^6^. Hypoxia not only alters tumor cell behavior but also directly affects infiltrating immune cells, including NK cells^7–10^. In addition, infectious environments present local hypoxia due to tissue damage and increased oxygen consumption^11^. Cellular adaptation to hypoxia is primarily mediated by a family of oxygen-sensitive transcription factors known as hypoxia-inducible factors (HIFs)^12^. The active HIF complex consists of an oxygen-regulated α-subunit (HIF-1α or HIF-2α) and a constitutively expressed β-subunit (ARNT). Under normoxic conditions, prolyl hydroxylases (PHDs) target HIF-α for proteasomal degradation, but in hypoxia, HIF-α is stabilized, dimerizes with HIF-β, and translocates to the nucleus to initiate transcription of hypoxia-responsive genes through binding to hypoxia response elements^13^. These genes regulate, amongst others, angiogenesis, metabolism, cell survival, and immune responses^14^.

While HIF-1α is expressed in most cell types, HIF-2α shows a more tissue-restricted pattern. Both isoforms shape immune-cell fate in hypoxia, but their exact functions in NK cells are only partly resolved. Most tumor-focused studies report that hypoxia dampens NK-cell cytotoxicity and that genetic or pharmacologic blockade of HIF-1α rescues degranulation, IFNγ release and target-cell killing^15,16^. Single-cell RNA-sequencing of human tumor infiltrates even classifies HIF-1α as an immune checkpoint that restrains NK cells^16^. However, some contrasting findings suggest that HIF-1α may also support NK cell function in specific contexts. For instance, during viral infection, HIF-1α is required for metabolic adaptation and survival of NK cells^17^. Similarly, in ex vivo-expanded NK cells primed with interleukin (IL)-15, HIF-1α supports NK cell effector functions even under hypoxic conditions^18^. Notably, IL-15 stimulation induces HIF-1α accumulation in NK cells also in normoxia^19^. Indeed, HIFs can also be stabilized by inflammatory signals, exceeding their classical roles as a hypoxia sensor. Altogether, these findings argue that HIF-1α can be either supportive or inhibitory in NK cells depending on the activation state, metabolic demand and hypoxic context. Far less is known about HIF-2α in NK cell biology. In murine NK cells, HIF-2α inhibition did not counteract the elevated effector responses of HIF1A knockout NK cells^16^. Intriguingly, insertion of a non-degradable EPAS1 transgene, encoding for HIF-2α, turned CD8-T cells into hyperfunctional, tumor-rejecting cells^20^. Hence, elucidation of the uncovered HIF-2α territory might hold potential for NK cell applications.

In this study, we comparatively looked into the contribution of HIF isoforms to primary human NK cell functioning. Since genetic modification remains technically challenging in primary human NK cells and could result into compensatory rewiring, we employed well-known small-molecule inhibitors targeting HIF-1α, HIF-2α, or PHDs to further dissect their contributions to NK cell biology. As the functional relevance of HIF-2α in NK cells remains largely unexplored, uncovering its role could reveal novel regulatory pathways and identify new targets to enhance NK cell-mediated therapies.

## Materials & Methods

### Ethics statement

Primary human NK cells were used, derived from buffy coats that were obtained from adult volunteer whole blood donations at the Blood Transfusion Center of the Red Cross-Flanders (Mechelen, Belgium). The use of these specimens was approved by the local Ethics Committee of the Antwerp University Hospital and the University of Antwerp (EC5488).

### Cell culture

Peripheral blood mononuclear cells were isolated from the buffy coats using Lymphoprep (7851, STEMCELL Technologies) density gradient centrifugation. Next, untouched primary human NK cells were isolated using negative magnetic activated cell sorting with the MojoSort Human NK Cell Isolation Kit (480054, Biolegend), according the manufacturer’s protocol. Following isolation, NK cells were cultured overnight in a standard incubator at 37 °C and 5 % CO_2_ in Roswell Park Memorial Institute (RPMI) 1640 medium (52400025, Life Technologies) supplemented with 10 % fetal bovine serum (FBS; 10270106, Life Technologies), 2 mM L-glutamine (25030024, Life Technologies) and 1 mM sodium pyruvate (11360070, Life Technologies) in. Addition of compounds and stimuli depended on the downstream assay.

The human chronic myelogenous leukemia K562 cell line is considered a golden standard target cell line for NK cell-mediated cytotoxicity due to its sensitivity. K562 cells were cultured in Roswell Park Memorial Institute (RPMI) 1640 medium (52400025, Life Technologies) supplemented with 10% FBS, 2mM L-glutamine, and 100 U/ml penicillin and 100 µg/ml streptomycin (15140122, Life Technologies) in a standard incubator at 37 °C and 5% CO_2_.

### NK cell-mediated cytotoxicity

Following NK cell isolation, the cells were cultured overnight at a density of 1.5×10^6^ cells/ml in the presence of 2 ng/ml IL-15 (130-095-765, Miltenyi Biotec) and compounds, i.e. 0.5% DMSO (D12345, Thermo Fisher Scientific) as vehicle control, 20 µM KC7F2 (S7946, Selleck Chemicals) to inhibit HIF-1α, 10 µM PT2385 (S8352, Selleck Chemicals) to inhibit HIF-2α, 10 µM KC7F2 plus 20 µM PT2385 to inhibit both HIF isoforms, or 10 µM MK-8617 (S8443, Selleck Chemicals) to inhibit PHDs. The next day, NK cells were washed to avoid the inhibitors influencing the target cells. Following cell counting on a TC20 cell counter (Biorad), taking viability into account by using a 1:1 dilution in Trypan blue (15250061, Life Technologies) and 200×10^3^ NK cells were plated per condition in a U-bottom 96-well plate (650180, Greiner Bio-One) to achieve a 5:1 effector:target ratio.

We used a flow cytometric cytotoxicity as described previously, with minor adjustments^21^. K562 target cells were pre-labeled with PKH67 using the PKH67 Green Fluorescent Cell Linker Mini Kit (MIDI67-1KT, Sigma-Aldrich), according to the manufacturer’s instructions, to allow specific gating on the target population. Per condition 40×10^3^ PKH67-labeled K562 cells were added. Tumor cells without NK cells served as controls. After 4 hours, the cells were washed and processed for analysis, dependent on the flow cytometer used. For analysis on a Quanteon (Novocyte), cells were washed with FACS buffer and subsequently stained with 2% 7-AAD (420403, Biolegend) and 1% AnnexinV-APC (550475, Biolegend) in 1X Annexin Staining Buffer (556454, BD Biosciences) for 10 min prior to flow cytometric acquisition. For analysis on a MACSQuant Analyzer 16 (Miltenyi Biotec) cells were washed with PBS buffer and subsequently stained for 20 min at room temperature with 2% LIVE/DEAD Fixable Aqua Dead Cell Stain (L34966, Life Technologies) in complete medium. Following a washing step with FACS buffer, a 10 min staining with 1% AnnexinV-APC on 1X Annexin Staining Buffer preceded flow cytometric acquisition. Live K562 cells were defined as 7-AAD-/AnnV-or LD-Aqua-/AnnV-cells of the PKH67^+^ population. NK cell-mediated cytotoxicity was calculated as: % killing = 100 % - (Live K562 cells with NK cells / Live K562 cells without NK cells) * 100.

### Intracellular cytokine staining

Following NK cell isolation, the cells were cultured overnight at a density of 1.5×10^6^ cells/ml in the presence of 2 ng/ml IL-15. The next day, NK cells were counted on a TC20 cell counter, taking viability into account, and 200×10^3^ NK cells were plated per condition in a U-bottom 96-well plate. Treatments were added to a concentration as described above. After 1 hour, 1X ionomycin Cell stimulation cocktail (00497093, Thermo Fisher Scientific) was added. One hour later Brefeldin A (00450651, Thermo Fisher Scientific) and Monensin (00450551, Thermo Fisher Scientific) were added to a 1X concentration. Cells were then incubated for another 3 hours. Then, cells were washed with PBS (14190250, and incubated with 2% LIVE/DEAD Near-IR Dead Cell Stain (L34976, Life Technologies) in medium for 20 min at room temperature. Next, cells were fixed and permeabilized using the FoxP3 / Transcription Factor Staining Kit (11500597, Thermo Fisher Scientific), according to the manufacturer’s protocol. Prior to staining, cells were blocked with 1:10 human serum (S1-LITER, Sigma-Aldrich) for 30 min. Intracellular staining was performed with 1:100 anti-human TNFα-BV605 (502935, Biolegend) 1:100 anti-human IFNγ-BV605 (502535, Biolegend), or 1: 100 anti-human Granzyme B (GZMB)-Alexa Fluor 6478 (515405, Biolegend). Samples were acquired on a MACSQuant Analyzer 16. Mean fluorescence of the positive cell population was determined and the MFI ratio of inhibitor versus vehicle was calculated for this population.

### Mitochondrial dyes staining

Following NK cell isolation, the 300×10^3^ NK cells were cultured overnight at a density of 1.5×10^6^ cells/ml in the presence of 20 ng/ml IL-15 and 100 IU/ml IL-2 (130-097-745, Miltenyi Biotec) to activate the NK cells. HIF-targeting compounds KC7F2 and PT2385 were added in the concentrations and conditions as described above. The next day, cells were washed with serum-free medium and resuspended in serum-free-medium containing 25 nM TMRM (M20036A, Thermo Fisher Scientific) or 1 µM MitoSOX Red (M36008, Thermo Fisher Scientific) to assess mitochondrial membrane potential and mitochondrial reactive oxygen species (mtROS), respectively. Following a 30 min incubation at 37 °C, cells were washed and stained with 2% LIVE/DEAD Near-IR Dead Cell Stain for 20 min at room temperature. Samples were acquired on a Novocyte Quanteon. Mean fluorescence of the positive cell population was determined and the MFI ratio of inhibitor versus vehicle was calculated for this population. Mean fluorescence of the positive cell population was determined and the MFI ratio of inhibitor versus vehicle was calculated for this population.

### Statistics

All experiments were performed using at least three independent donors in at least two independent experiments. Flow cytometric analyses were performed using FlowJo v10.10.0 (FlowJo LLC). Graphical visualization and statistical analyses were performed using GraphPad Prism v10.5.0 (Dotmatics). Normality was not assumed. Non-parametric paired testing was performed. To compare two groups, Wilcoxon matched-pairs signed-rank test was used. For comparisons involving more than two groups, Friedman test with Dunn’s correction for multiple comparisons was performed. A p-value < 0.05 was considered statistically significant. A p-value < 0.10 was considered a statistical trend.

## Results

### HIF-1α but not HIF-2α inhibition impairs NK cell-mediated killing in normoxia

Since the primary function of NK cells towards tumor cells is killing of the latter, we first assessed the role of HIF isoforms in the cytotoxic function of NK cells under normoxic conditions. NK cells were overnight cultured with recognized inhibitors against HIF-1α (KC7F2), HIF-2α (PT2385), or both. The next day the inhibitors were washed away prior to the coculture, in order to avoid them affecting the tumor cells directly. HIF-1α-inhibited NK cells displayed a profound reduced capacity to kill K562 leukemia cells (Fig. 1). On the other hand, treatment with PT2385 to inhibit HIF-2α did not impact the killing capacity, generating similar effects as vehicle-treated NK cells. When both KC7F2 and PT2385 were added in order to shut off both isoforms, similar results were obtained to HIF-1α inhibition, with statistical trends versus vehicle and PT2385. Overall, these results indicate that HIF-1α is involved in the cytolytic machinery of NK cells, while HIF-2α is redundant for this function.

**Figure 1.**
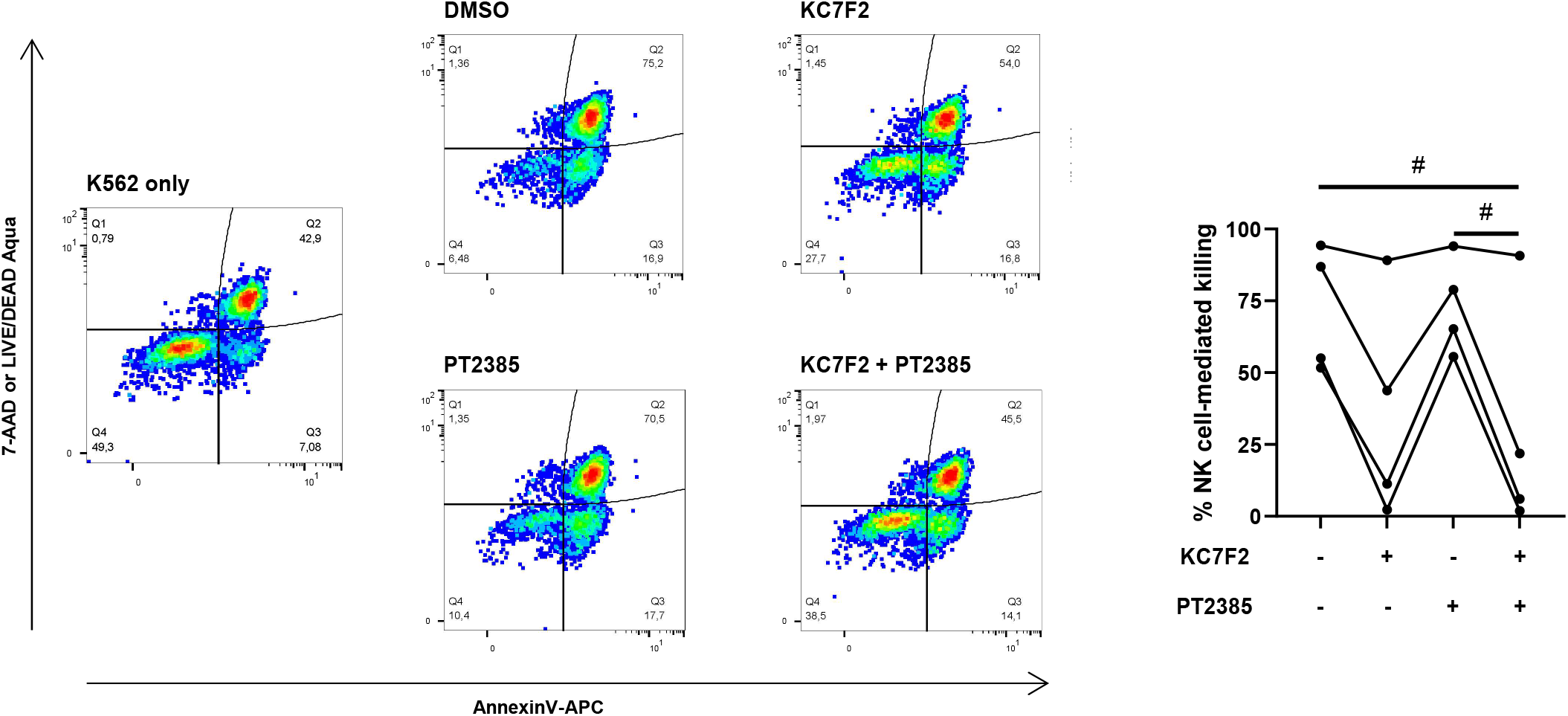
Inhibition of HIF-1α impairs NK cell-mediated cytotoxicity. NK cells treated with vehicle (DMSO), HIF-1α inhibitor KC7F2, HIF-2α inhibitor PT2385, or both HIF inhibitors combined, were cocultured with PKH67-labeled K562 cells for 4 hours (n = 4). Cytotoxicity was determined based on flow cytometry measurements. Left: representative flow cytometry dot plots. Friedman test with Dunn’s correction for multiple comparisons. # p < 0.10. HIF: hypoxia-inducible factor; NK cell: natural killer cell.

### HIF isoforms differently influence cytokine production, not granzyme B

NK cells are equipped with a set of effector tools to mediate the killing of tumor cells. Their primary mode of action regard perforin and granzyme-mediated cytotoxicity. Given the reduction in tumor cell killing when exposed to the HIF-1α inhibitor KC7F2, we investigated its effect on GZMB production using a stimulation assay in the presence of the inhibitors. For any condition, nearly all NK cells possessed GZMB, as physiologically expected (Fig. 2A). In addition, the amount of GZMB did not change either (Fig. 2A, MFI ratio, and Suppl. Fig. S1A). Looking further at TNFα and IFNγ – intracellular cytokines with the potential to kill cancer cells though predominantly acting in an immunoregulatory role – patterns similar to the killing assay were observed (Fig. 2B-C and Suppl. Fig. S1B-C). Compared to control, significantly less NK cells were positive for these cytokines when HIF-1α was inhibited. In addition, the positive cells also produced less of the respective cytokine, as indicated by the MFI. In contrast again, HIF-2α inhibition did not appear to have a consistent effect on TNFα. PT2385 reduced both the frequency of IFNγ^+^ NK cells and their IFNγ production, although statistically non-significant. Combined inhibition of both isoforms yielded results comparable to only HIF-1α inhibition. Altogether, GZMB content appears unaffected by HIFs, hence not probable to cause the observed reduction in killing capacity upon HIF-1α inhibition. However, cytokine production is vastly impacted by HIF-1α, while HIF-2α seems to play a minor role at best.

**Figure 2.**
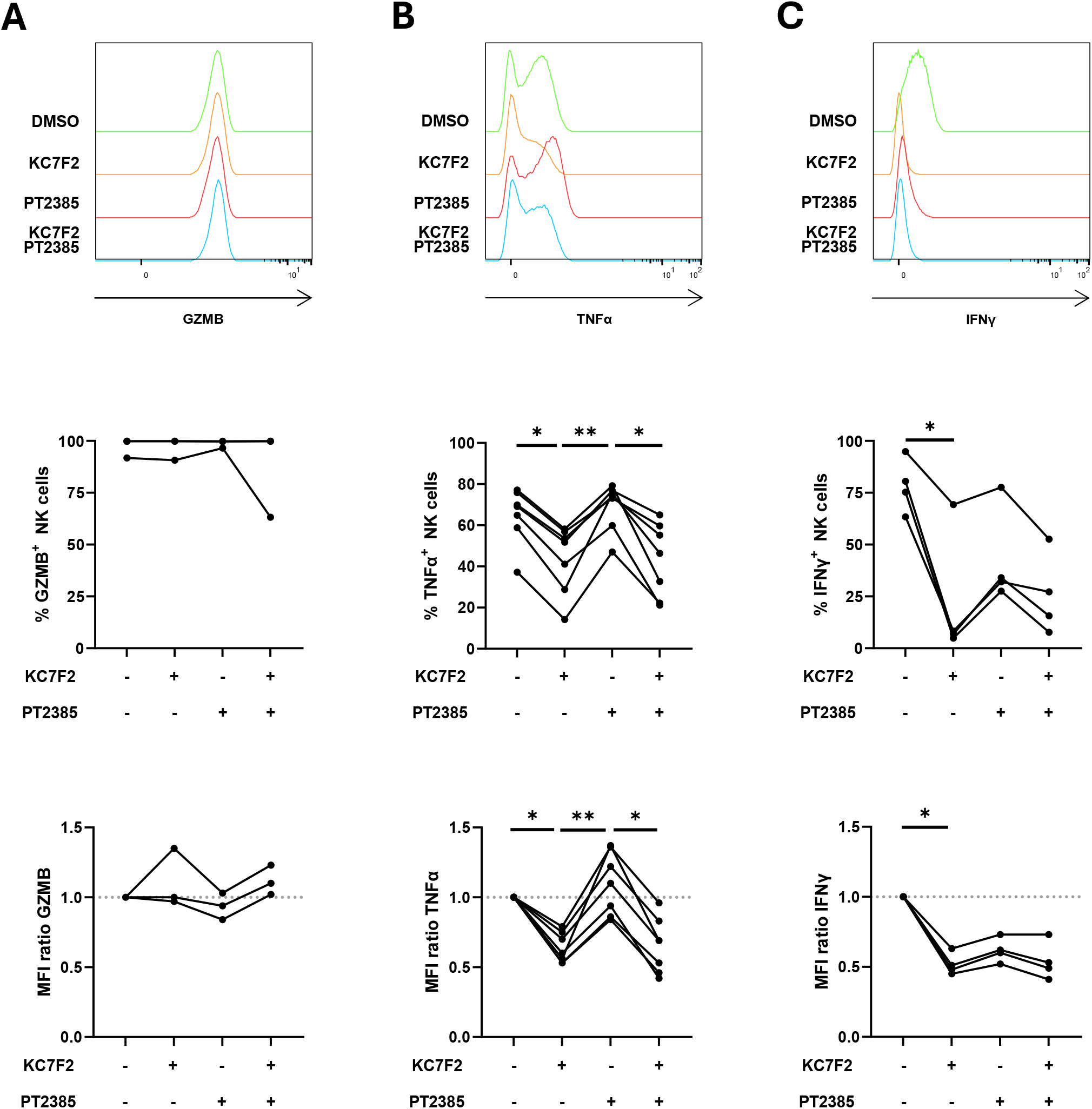
Inhibition of HIF-1α impairs pro-inflammatory cytokine production. NK cells were stimulated with PMA/ionomycin on treatment with vehicle (DMSO), HIF-1α inhibitor KC7F2, HIF-2α inhibitor PT2385, or both HIF inhibitors combined. Intracellular cytyokine staining of (A) granzyme B (GZMB), (B) TNFα, or (C) IFNγ was evaluated by flow cytometry (n=3-7). Top row: representative flow cytometry histograms. Middle row: percentage positive NK cells. Bottom row: ratio of MFI HIF inhibition condition over MFI vehicle condition. Friedman test with Dunn’s correction for multiple comparisons. * p < 0.05, ** p < 0.01. HIF: hypoxia-inducible factor; MFI: mean fluorescence intensity; NK cell: natural killer cell.

### Inhibition of HIF-1α increases mitochondrial ROS in NK cells

Among their pleiotropic functions, HIF mediate metabolic adaptations. Since immunometabolism is key to effective NK cell functioning, we looked at mitochondrial readouts following overnight stimulation with IL-15 and IL-2. Inhibition of HIF-1α significantly increased the number of mtROS^+^ NK cells, as well as the mtROS content in those cells, as indicated by MitoSOX staining (Fig. 3A and Suppl. Fig. S1D). In concordance with the other parameters, this effect was not observed with HIF-2α inhibition, yet replicated when both HIF isoforms were targeted. In addition to mtROS, we also analyzed the mitochondrial membrane potential, which was not consistently affected in any of the HIF inhibitory conditions (Fig. 3B and Suppl. Fig. S1E).

**Figure 3.**
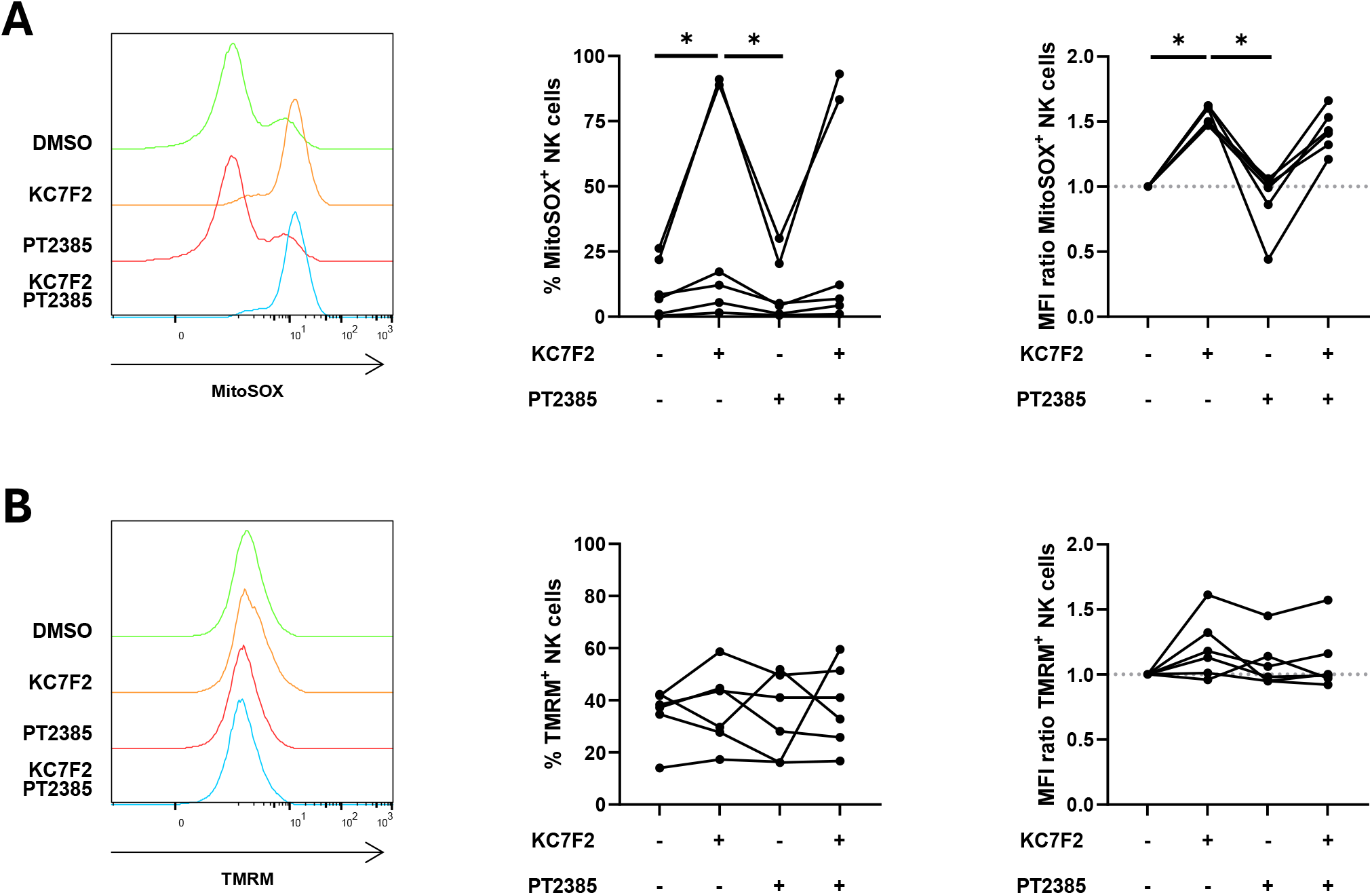
Inhibition of HIF-1α increases mitochondrial ROS. NK cells were overnight stimulated with IL-15 and IL-2 and treated with vehicle (DMSO), HIF-1α inhibitor KC7F2, HIF-2α inhibitor PT2385, or both HIF inhibitors combined. Mitochondrial parameters were flow cytometrically assessed via (A) MitoSOX Red voor mtROS, and (B) TMRM for mitochondrial membrane potential (n=6). Left: representative flow cytometry histograms. Middle: percentage positive NK cells. Right: ratio of MFI HIF inhibition condition over MFI vehicle condition. Friedman test with Dunn’s correction for multiple comparisons. * p < 0.05. HIF: hypoxia-inducible factor; MFI: mean fluorescence intensity; mtROS: mitochondrial ROS; NK cell: natural killer cell.

### HIF stabilization does not stimulate NK cell effector functioning

Given that we observed that inhibition of HIF-1α mitigates effector functions and metabolism of NK cells, we hypothesized that stabilization of HIF-1α would stimulate those aspects of NK cell functioning. Therefore, we repeated the different assays with a pan-PHD inhibitor, MK8617. Since PHD prime HIF-α isoforms for degradation, we would expect that this approach would result in heightening of both HIF-1α and HIF-2α Surprisingly, HIF stabilization did not result in increased effector functioning and potential, nor consistent metabolic alterations (Fig. 4 and Suppl. Fig. 2). This suggest that while HIF-1α inhibition impairs effector functions, its stabilization is not sufficient to stimulate these same functions, at least in circumstances investigated.

**Figure 4.**
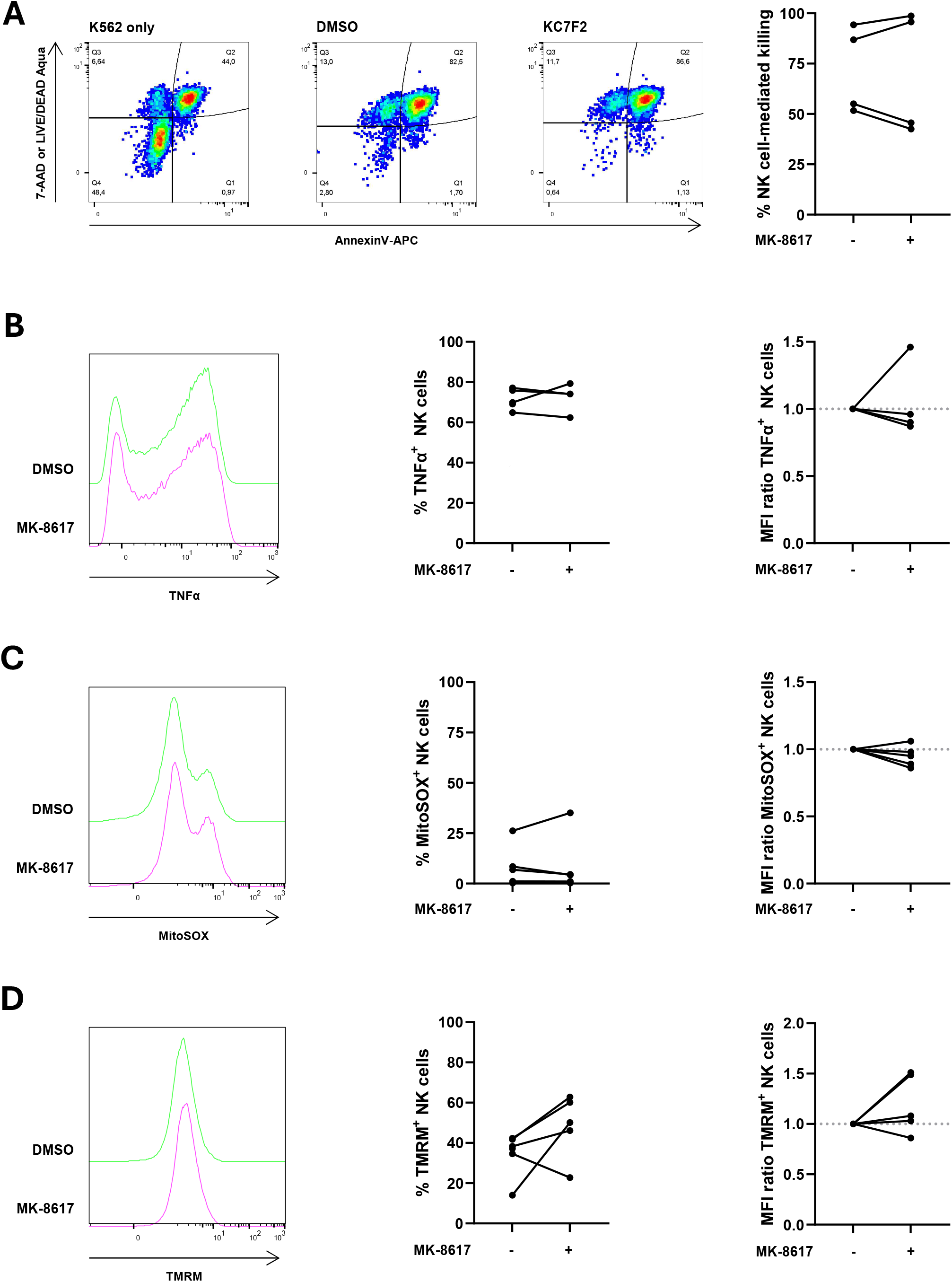
HIF stabilisation does not affect NK cell effector function. NK cells were treated with vehicle (DMSO) or PHD inhibitor MK-8617 and analyzed for (A) NK cell-mediated cytotoxicity, (B) pro-inflammatory cytokine production, (C) mtROS, and (D) mitochondrial membrane potential (n=4). Wilcoxon pairs signed-rank test. HIF: hypoxia-inducible factor; MFI: mean fluorescence intensity; mtROS: mitochondrial ROS; NK cell: natural killer cell; PHD: prolyl hydroxylase.

## Discussion

In this study we investigated the contribution of HIF-1α and HIF-2α to primary human NK cell effector functions in normoxia through pharmacological inhibition. Our results indicate that under normoxic conditions HIF-1α plays a dominant role in cytotoxicity and pro-inflammatory cytokine production upon activation. GZMB remained stable in HIF-1α-inhibited human NK cells, although in contrast to murine HIF1A^KO^ NK cells^22^. Together with increased mitochondrial ROS production, this could indicate HIF-1α rather works through transcriptional and metabolic adaptations rather than impacting the cytolytic machinery in toto, in line with observation of Coulibaly et al. in a IL-15 context^19^. In contrast to HIF-2α which did not appear to participate to a large extent.

Our observation that HIF-1α is involved in effector functioning of NK cells is in line with the reported, although contrasting literature. In support of our findings, decreased degranulation and IFNγ production of HIF1A^KO^ murine NK cells directly ex vivo was reported, indicative for impairment tumor cell killing^22,23^. In contrast, Ni et al. reported increased killing capacity by human NK cells treated with HIF-1α inhibitor KC7F2 for 7 days^16^. HIF1A^KO^ in 7 days-activated human NK cells, followed by a 4-days expansion protocol corroborated this by detecting strengthened killing and TNF gene expression^15^. Notably, Kennedy et al. demonstrated that HIF1A^KO^ did not affect the killing capacity of expanded NK cells, suggesting that that expanded NK cells are primed for hyperfunctioning^8^. The discrepancy in the role of HIF-1α might be contextual. In acute and short-term situations, HIF-1α could help NK cells to maintain and execute their function, through HIF-1α-driven glycolysis and preserved effector functions. Our mitochondrial analysis favored mitochondrial metabolism upon HIF-1α inhibition, supporting that HIF-1α drives glycolytic phenotype, required for effector functioning^24^. However, chronic activation due to chronic inflammatory signals or prolonged hypoxia, might lead to NK cell adaptation or exhaustion, and eventually a functional decline^8,25^. Particularly in cancer this could be an important nuance to take into account, since hypoxia is a characteristic of many solid tumors. Interestingly, while Krzywinska et al. observed impaired tumor cell killing by HIF1A^KO^ NK cells in vitro, tumor reduction was observed in vivo, attributed to stimulation of non-productive angiogenesis due to reduced soluble VEGFR-1 expression by HIF1A^KO^ NK cells^23^. Overall, while the involvement of HIF-1α in NK cell effector functions is undoubted and holds potential therapeutic application, its duality warrants further investigation to elucidate its contextual dependence.

In contrast to HIF-1α, HIF-2α did not participate in NK cell effector functions in our design, including no counteraction of the HIF-1α phenotype in combinatorial targeting. This corroborates a previous study stating HIF-2α does not antagonize HIF-1α in NK cell effector response, although this response was opposite to what we observed^16^. We only cultured the NK cells short-term with the inhibitor, while HIF-2α has slower stabilization dynamics compared to HIF-1α^26^. While Ni et al. did not find HIF-2α inhibition to counteract HIF-1α after 7 days of culture, they actually observed improved IFNγ production when HIF-2α was inhibited^16^. This supports the engineering strategy Velica et al., who demonstrated that CD8^+^ T cells with ectopically stabilized HIF-2α featured enhanced anti-tumor functions^20^. Contrasting though, this did not involve increased IFNγ expression. Hence, while current studies indicate a limited role of HIF-2α in short-term NK cell activation, further research on longer periods is warranted.

In conclusion, our study indicates that in a short-time window, inhibition of HIF-1α disrupts NK cell effector functioning, suggesting that HIF-1α contributes to NK cell duties in acute contexts. On the other hand, HIF-2α was not implicated in NK cell functioning during this time window. Overall, HIF isoforms require a deeper and contextual understanding regarding their roles in NK cell functioning in order to move forward with these targets towards therapeutic application.

## Acknowledgements

This work was supported by Research Foundation Flanders (G040120N; J.M. 11PHM24N) and by the University of Antwerp (BOF and IOF grant numbers to J.D.W. FFB230049, FFB230316, FFI230209, and J.d.B. FFB240247). T.V. received support from Kom op tegen Kanker (Stand up to Cancer), the Flemish cancer society (grant number OZ10180). J.D.W. received support from the ME TO YOU foundation (grant number OZ8546). Part of the research was supported by donations from different donors, including Willy Floren, Dedert Schilde vzw and the Vereycken family.

## Notes

### Competing Interest Statement

The authors have declared no competing interest.

