## Supplemental Figure 1 for "HIF isoforms contribute distinctly during human NK cell activation"

### Supplementary Figure S1

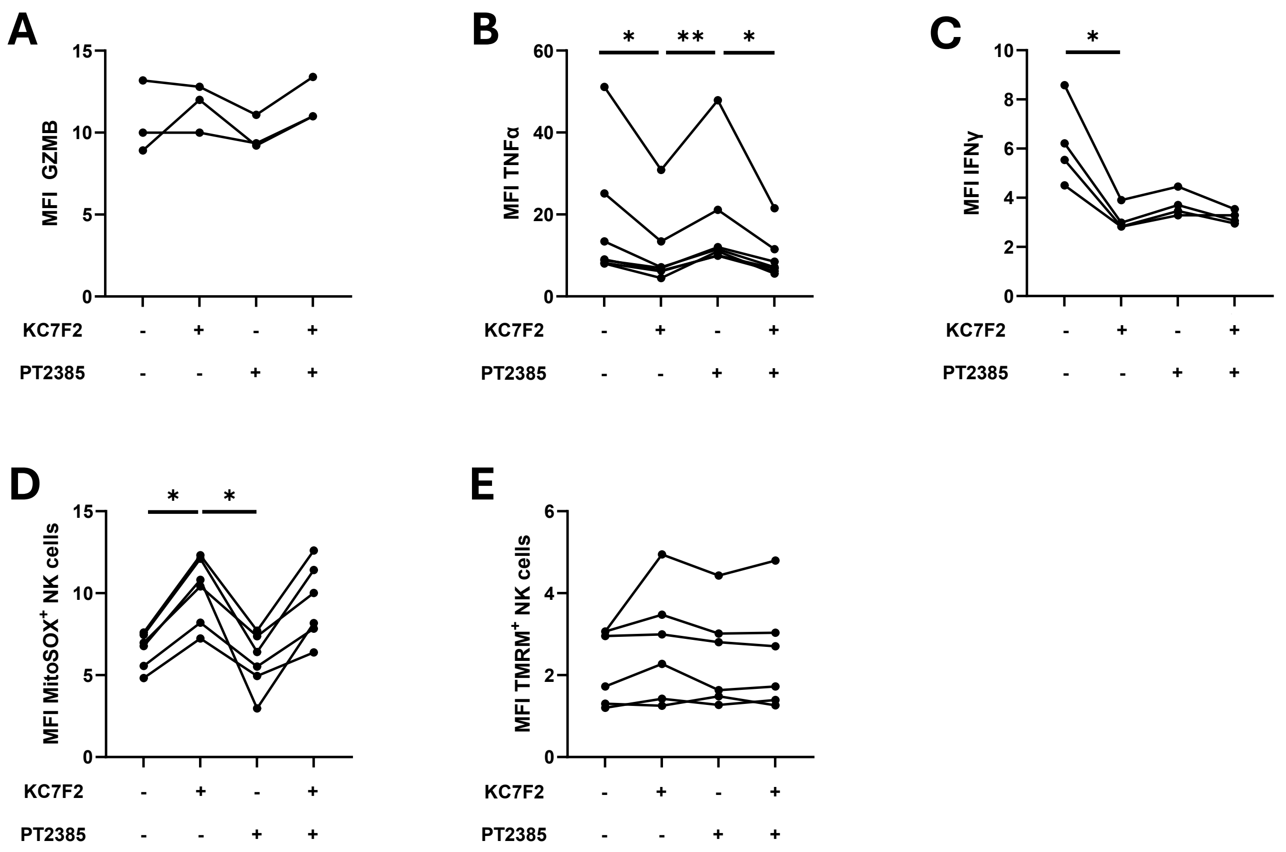

**Supplementary Figure S1. Raw MFI data of HIF inhibition conditions.** MFI of NK cells expressing (A) GZMB, (B) TNFα, or (C) IFNγ as well as those positive for (D) MitoSOX, and (E) TMRM dyes (n=3) nK cells were treated with vehicle (DMSO), HIF-1α inhibitor KC7F2, HIF-2α inhibitor PT2385, or both HIF inhibitors combined. Friedman test with Dunn's correction for multiple comparisons. * $p < 0.05$, ** $p < 0.01$. GZMB: granzyme B; HIF: hypoxia-inducible factor; MFI: mean fluorescence intensity; NK cell: natural killer cell
