## Supplemental Figure 2 for "HIF isoforms contribute distinctly during human NK cell activation"

### Supplementary Figure S2

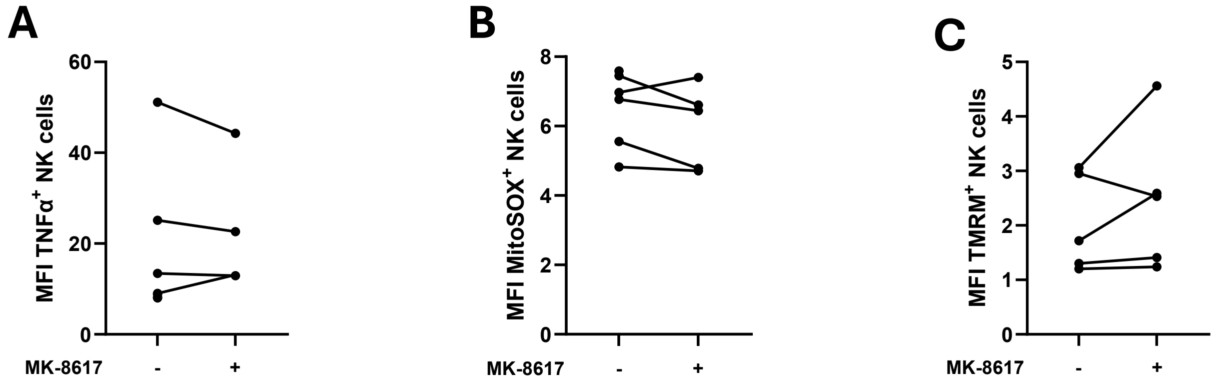

**Supplementary Figure S2. Raw MFI data of HIF stabilisation condition.** MFI of NK cells expressing (A) TNFα, or being those positive for (B) MitoSOX, and (C) TMRM dyes (n=3° nK cells were treated with vehicle (DMSO) or PHD inhibitor MK-8617 Wilcoxon pairs signed-rank test. * $p < 0.05$, ** $p < 0.01$. HIF: hypoxia-inducible factor; MFI: mean fluorescence intensity; NK cell: natural killer cell
